# KinasepKipred: A Predictive Model for Estimating Ligand-Kinase Inhibitor Constant (*pK_i_*)

**DOI:** 10.1101/798561

**Authors:** KC Govinda, Md Mahmudulla Hassan, Suman Sirimulla

**Affiliations:** Computational Science Program, The University of Texas at El Paso, Texas, USA; Department of Computer Science, The University of Texas at El Paso, Texas, USA; Department of Pharmaceutical Sciences, School of Pharmacy, The University of Texas at El Paso, Texas, USA

## Abstract

Kinases are one of the most important classes of drug targets for therapeutic use. Algorithms that can accurately predict the drug-kinase inhibitor constant (*pK_i_*) of kinases can considerably accelerate the drug discovery process. In this study, we have developed computational models, leveraging machine learning techniques, to predict ligand-kinase (*pK_i_*) values. Kinase-ligand inhibitor constant (*K_i_*) data was retrieved from Drug Target Commons (DTC) and Metz databases. Machine learning models were developed based on structural and physicochemical features of the protein and, topological pharmacophore atomic triplets fingerprints of the ligands. Three machine learning models [random forest (RFR), extreme gradient boosting (XGBoost) and artificial neural network (ANN)] were tested for model development. The performance of our models were evaluated using several metrics with 95% confidence interval. RFR model was finally selected based on the evaluation metrics on test datasets and used for web implementation. The best and selected model achieved a Pearson correlation coefficient (R) of 0.887 (0.881, 0.893), root-mean-square error (RMSE) of 0.475 (0.465, 0.486), Concordance index (Con. Index) of 0.854 (0.851, 0.858), and an area under the curve of receiver operating characteristic curve (AUC-ROC) of 0.957 (0.954, 0.960) during the internal 5-fold cross validation.

**Availability:** GitHub: https://github.com/sirimullalab/KinasepKipred, Docker: `sirimullalab/kinasepkipred`

**Implementation:** https://drugdiscovery.utep.edu/pki/

**Graphical TOC Entry:** 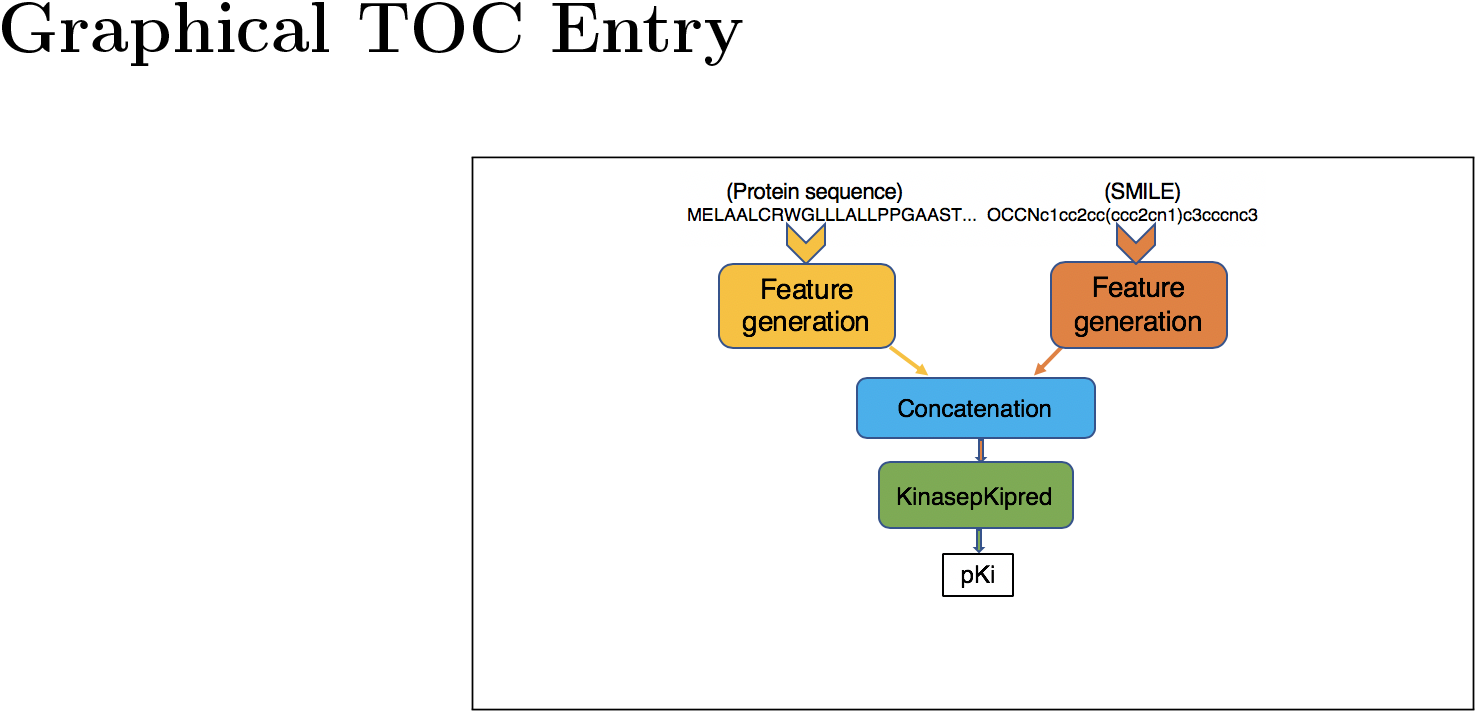

## Introduction

The interaction between drug and target facilitates the drug side effect prediction,^1^ drug repurposing^2^ and many others. Biochemical experiment methods for drug target interactions are found highly costly and take a lot of time,^3^ whereas computational methods are efficient, faster and more convenient.^4^ Proteins are the good targets in drug design^5^ and get activated or inhibited by drug compounds. The binding of protein with other ligand is found to be specific and it plays crucial role in many biological functions.^6^ Protein kinase is an enzyme that plays a major role in the signal transduction of cells by transferring a phosphate group from adenosine triphosphate (ATP) to other proteins to their serine, threonine or tyrosine residues.^7^ In this study, we are focused on protein kinases because of their importance as drug targets for therapeutic use. They play important roles in a wide range of diseases such as cardiovascular disorders, inflammatory diseases, gastrointestinal stromal tumors and cancer, and can serve as drug targets for therapeutic use.^8^ Kinase inhibitors that inhibit the activity of deregulated protein kinases very efficacious in treating many diseases.^9^ The US food and drug administration has approved 48 kinase inhibitors^9^ and the largest class of new drugs approved is for cancer treatment. The interaction between the kinase and ligand is usually measured as binding affinity values in terms of dissociation constant (*K_d_*), inhibition constant (*K_i_*) and half-maximal inhibitory concentration (*IC*_50_). In this paper, we have used only the data corresponding to (*K_i_*) values for our dataset. Algorithms that predict drug-target associations^10,11^ and binding affinities were previously published by other researchers. ^6,10,12–14^ Yamanishi et al.^10^ proposed a supervised machine learning approach for categorizing drug-target pairs as interacting or non-interacting based on an integrated model of chemical and genomic molecular profile^10^ as a binary class problem. Pahikkala et al.^12^ introduced a method KronRLS as a very first one to predict the non-binary drug target binding affinity values. After that, Simboost,^13^ DeepDTA^14^ and Indra et al.^6^ are the most recent studies based on machine learning models to predict the binding affinity scores. Simboost purposed the gradient boosting machine learning model by using feature engineering to predict the binding affinity values. It is the first non-linear method for continuous drug-target interaction prediction. DeepDTA is a deep learning based approach to predict drug-target binding affinity using only sequences of proteins and drugs. They used convolution neural networks (CNNs) to learn representations from the raw sequence data of proteins and drugs and fully connected layers for the prediction. Simboost and DeepDTA are based on Davis,^15^ Metz^16^ and KIBA^13^ data sets. Indra et al.^6^ developed a Random forest ML based on pdbbind database 2015. All aforementioned studies were generalized for all types of proteins and were not kinase specific and also purposed models were not evaluated by independent data sets. To our best knowledge, there are no algorithms that are specific to kinase *K_i_* predictions. In this study, the performance of three different machine learning algorithms Random forest (RFR), Extreme gradient boosting (XGBoost), and Artificial neural network (ANN) was developed and compared based on test dataset and external dataset. The best model based on evaluated statistical metrics is selected and used for web implementation.

## Methods

### Datasets

Data was obtained from two different databases 1) Drug Target Commons (DTC) ^17^and 2) Metz.^16^ DTC data contained 5,867,349 ligand-target pair associations. The dataset was populated by the bioctivity types *K_i_* (inhibition constant), *K_d_* (dissociation constant), and *IC*_50_ (half maximal inhibitory constant) for most of the ligand-target pairs. The “potent targets” and “potent inhibitors” were defined based on cut-offs for the four most popular bioactivity types (*K_d_*, *K_i_*, *IC*_50_, and activity). Cutoffs of ≤ 100 nM (i.e., ≥ *pK_i_* of 7) for the dose-response measurements (*K_d_*, *K_i_*, *IC*_50_) in biochemical assays, and ≤ 1000 nM (i.e., ≥ *pK_i_* of 6) for the dose-response measurements (*K_d_*, *K_i_*, *IC*_50_) in cell-based and other assay types^17^ were used. The median of the bioactivity values was taken where there were multiple bioactivity values. Since we were only focused on kinases with *K_i_*, we obtained only data pertinent to kinases with *K_i_* values. We found 67,894 instances (ligand-target pairs) from 5,983 compounds and 118 kinases. All *K_i_* values were converted into molar units and then recorded as the negative decadic logarithm as shown in equation (1). We used 75% of the data for the training set and 25% for the test set. The data set by Metz et al.^16^ was also used as an external evaluation for the model. It contained 150,000 instances of ligand-kinase *K_i_* values composed of more than 3,800 compounds tested against 172 protein kinases. We filtered out this Metz data to create a blind data set having 148 kinases with 240 compounds contributing 17,258 drug kinase pairs which are distinct from the DTC dataset and were not used in our training and test data.
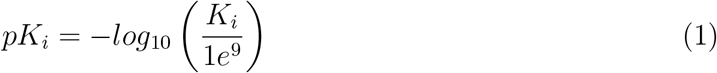

**Table 1:** Summary of datasets

| Data Sets | Kinases | Compounds | Interactions |
| --- | --- | --- | --- |
| DTC | 118 | 5983 | 67894 |
| Metz | 148 | 240 | 17258 |

**Figure 1:**
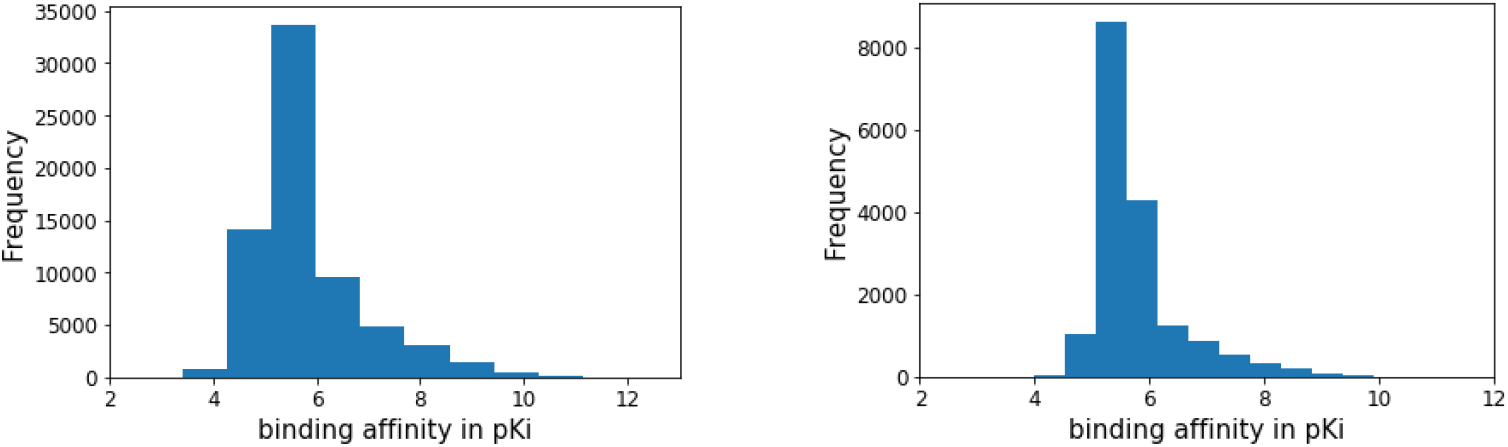
Distribution of datasets (a) Test dataset (b) Metz dataset

### Molecular features

#### Protein features

Propy,^18^ a protein feature generator algorithm was used to generate different features. The features that were used are as follows: *i*) Amino acid composition (20) *ii*) Dipeptide composition (400) *iii*) Moreau-Broto autocorrelation (240) *iv*) Moran autocorrelation (240) *v*) Geary autocorrelation (240) *vi*) composition transition distribution descriptors (147), *vii*) Type I pseudo amino acid composition descriptors (PACC) (30), *viii*) amphiphilic (Type II) pseudo amino acid composition descriptors (40) *ix*) sequence order coupling number (90) *x*) quasi sequence order descriptors (100) *xi*), and conjoint triad features (512).

#### Ligand features

MayaChemTools,^19^ a cheminformatic toolkit, was used to generate ligand features. SMILES, which represent ligands, were passed through a Perl script^20^ to generate topological pharmacophore atomic triplets fingerprints (TPATFP).^21–25^ The descriptive values were obtained in csv files containing fingerprint vector strings corresponding to molecular fingerprints. Pharmacophore atom types based on specified values were assigned to all nonhydrogen atoms of each molecule and a distance matrix was generated. A binned matrix was generated based on minimum distance, maximum distance and bin size values. The minimum distance (*D_min_*) and maximum distance (*D_max_*) corresponds to the number of bonds between two atoms. Pharmacophore atom triplets basis sets were generated for all unique atom triplets constituting atom pairs binned distance between the minimum distance and maximum distance. Atom triplets basis set were generated based on the following constraints (*i*) triangle rule, i.e., the length of each side of a triangle cannot exceed the sum of the lengths of the other two sides; and (*ii*) elimination of redundant pharmacophores related by symmetry. The possible values for pharmacophore atom types are: Ar, CA, H, HBA, HBD, Hal, NI, PI, RA. However, we have used the default values: HBD, HBA, PI, NI H, Ar. See Ref,^25^ where HBD= Hydrogen Bond Donor, HBA=hydrogen bond acceptor, PI=positively Ionizable, NI=negatively ionizable, Ar=aromatic, Hal=halogen, RA=ring atom, H=hydrophobic, and CA=chain atom. We have used the default values for *D_min_*(= 1), *D_max_*(= 10) and *B_size_*(= 2) and the number of distance bins *N_bins_*(= 5). Thus, our atom triplet basis set was of size 2692.

#### Model development

Models were developed mainly based on the grid search method with 5-fold cross validation. They were based on the Scikit-Learn machine learning library for Python. ^26^ We used the 25% of data for the test set and 75% of the data for the training set. Three different machine learning models (random forest, extreme gradient boosting, and artificial neural network) were developed and we compared their performances using several evaluation metrics. More detailed explanation of these three models are available in the supporting information. We also compared grid search vs random search^27^ for our random forest model to estimate the efficiency of each method.

## Results and Discussions

The developed models were evaluated using several metrics such as root-mean-square-error (RMSE), Pearson correlation coefficient (R), Spearman correlation coefficient (*ρ*), concor-dance index (Con. Index), and area under the receiver operating characteristic curve (AUC-ROC). The table 2,3 and 4 show the scores obtained for the test and Metz data set (external test dataset) using three models.

**Figure 2:**
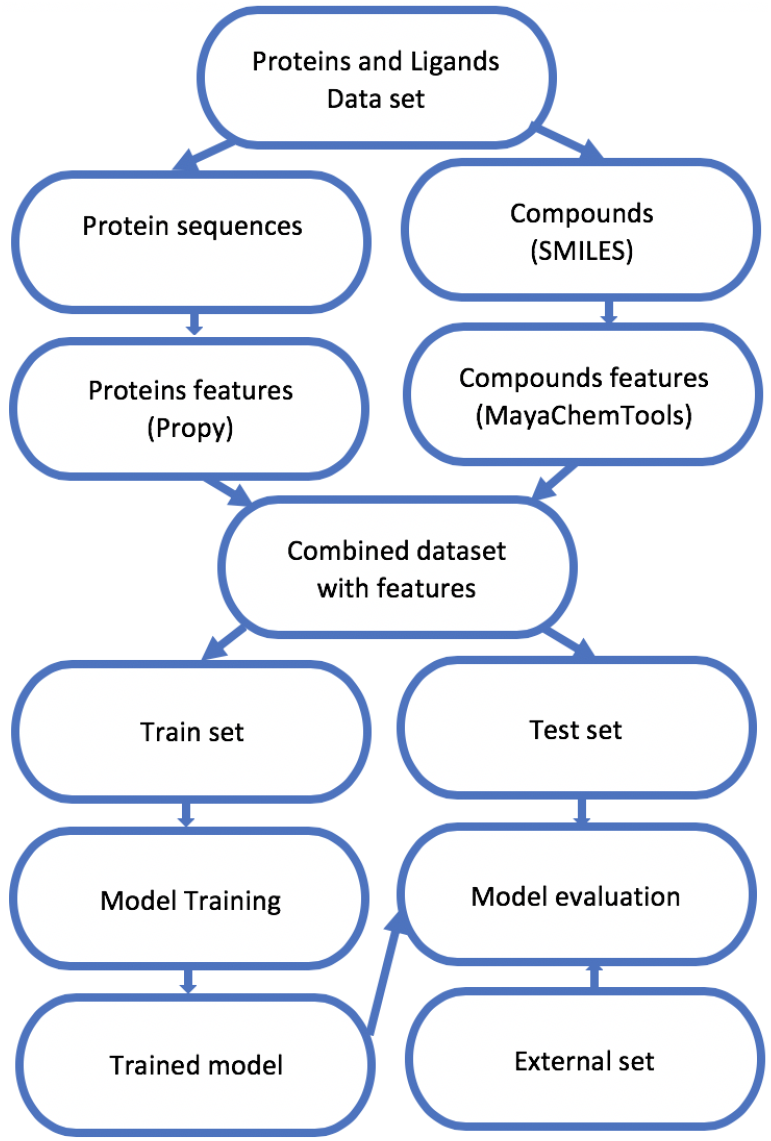
Project work flow

**Table 2:**
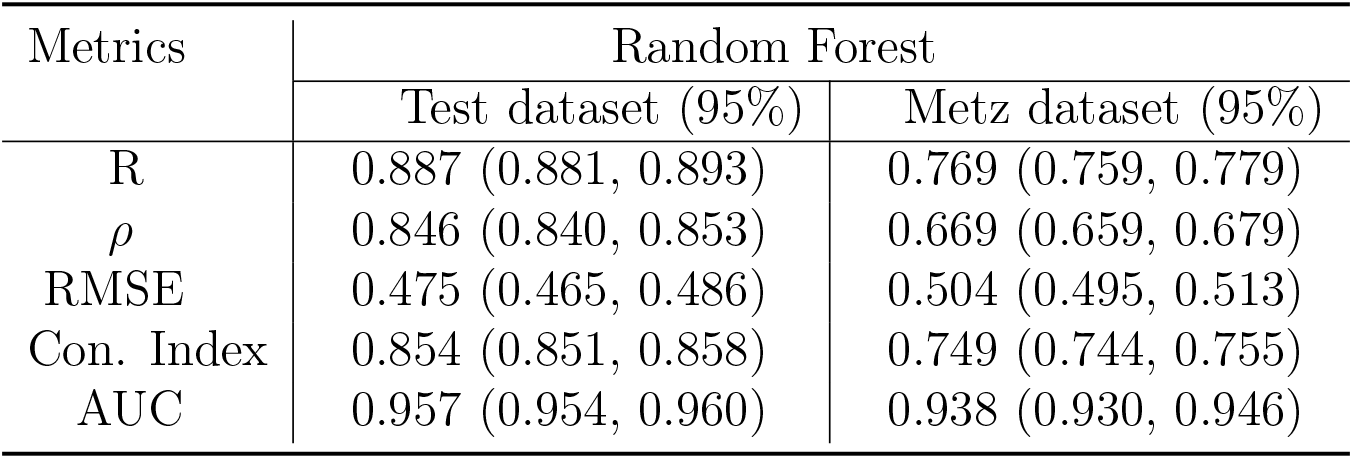
Performance of RF model on the test and Metz datasets

**Table 3:** Performance of XGBoost on the test and Metz datasets

| Metrics | XGBoost |  |
| --- | --- | --- |
|  | Test dataset (95%) | Metz dataset (95%) |
| R | 0.862 (0.855, 0.868) | 0.667 (0.656, 0.679) |
| $\rho$ | 0.795 (0.788, 0.802) | 0.574(0.563, 0.585) |
| RMSE | 0.522 (0.512, 0.533) | 0.582 (0.574, 0.591) |
| Con. Index | 0.805 (0.801, 0.810) | 0.703 (0.698, 0.709) |
| AUC | 0.946 (0.943, 0.950) | 0.922 (0.913, 0.931) |

**Table 4:** Performance of ANN on the test and Metz datasets

| Metrics | Artificial neural network |  |
| --- | --- | --- |
|  | Test dataset (95%) | Metz dataset (95%) |
| R | 0.798 (0.794, 0.803) | 0.60 (0.592, 0.608) |
| $\rho$ | 0.716 (0.710, 0.722) | 0.498 (0.489, 0.507) |
| RMSE | 0.631 (0.623, 0.639) | 0.619 (0.612, 0.627) |
| Con. Index | 0.779 (0.774, 0.784) | 0.658 (0.651, 0.665) |
| AUC | 0.933 (0.926, 0.940) | 0.887 (0.880, 0.894) |

Among three different models, random forest was found performing best with R 0.887, *ρ* 0.846, RMSE 0.475, Con. Index 0.854, and AUC 0.957 for the test data set and 0.769, 0.669, 0.503, 0.749, and 0.938 respective scores for the external data set. More details about the results and a comparative study can be found in the supporting information.

### Performance of different ML algorithms on the test dataset

Out of 67894 instances, 75% of the data was used for training with 5-fold cross validation and 25% of the data were used as a blind test set. The results obtained for the test set, for RFR, were R 0.887 (0.881, 0.893), *ρ* 0.846 (0.840, 0.853) RMSE 0.475 (0.465, 0.486), Con. Index 0.854(0.851, 0.858), and AUC 0.957 (0.954, 0.960); for XGBoost were R 0.862(0.855,0.868), ρ 0.795(0.788,0.802), RMSE 0.522(0.512,0.533), Con. Index 0.805(0.801,0.810), and AUC 0.946(0.943,0.950); and for ANN were R 0.798 (0.794, 0.803), *ρ* 0.716(0.710,0.722) RMSE 0.631(0.623,0.639), Con. Index 0.779 (0.774, 0.784), and AUC 0.933(0.926,0.940). All the metrics were computed with a 95% confidence interval. AUC is generally used for classification problems. Since this was a regression problem, quantitative values were converted into binary values using certain thresholds. We used 10 different equally spaced thresholds lying between 6 and 8 (DTC^17^) and the average AUC was calculated. After comparing the results for all the three models, we found that RFR outperformed XGBoost and ANN. XGboost had slightly lower performance than RFR, but it was better than ANN. ANN did not work well with our dataset.

**Figure 3:**
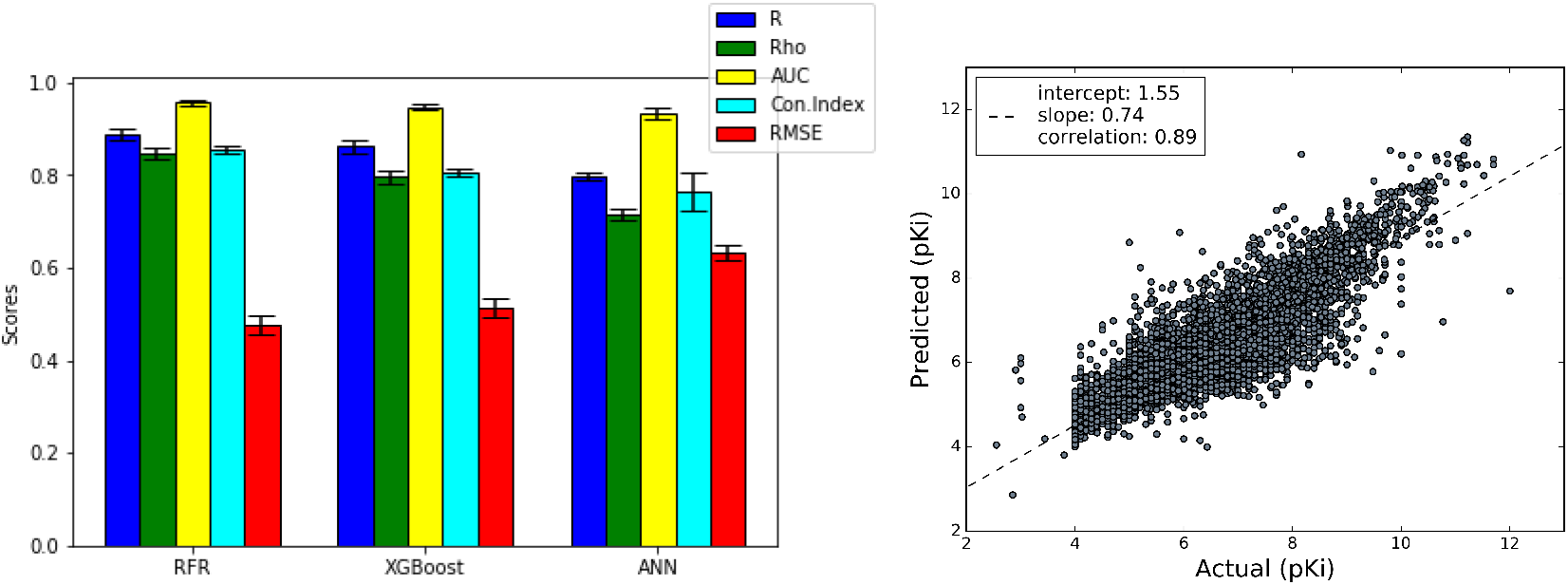
(a) Comparison of models on the test dataset (b) Predictions from Random Forest against actual values on the test dataset.

### Performance of different ML algorithms on an external dataset

All the three models were further tested on the Metz dataset.^16^ The results obtained for RFR were R 0.769 (0.759, 0.779), *ρ* 0.669 (0.659, 0.679), RMSE 0.504 (0.495, 0.513), Concordance Index 0.749 (0.744, 0.755), and AUC 0.938 (0.930, 0.946). For XGBoost, R 0.667 (0.656, 0.679), *ρ* 0.574 (0.563, 0.585), RMSE 0.582 (0.574, 0.591), Concordance Index 0.703 (0.698, 0.709), and AUC 0.922 (0.913, 0.931) and and for ANN, R 0.60 (0.592, 0.608), *ρ* 0.498 (0.489, 0.507), RMSE 0.619(0.612,0.627), Con. Index 0.658 (0.651, 0.665), and AUC 0.887 (0.880, 0.894). All the metrics were computed with a 95% confidence interval. Model performance on an external set was somewhat low compared to the model performance on the test set, but AUC showed a promising result for RFR and XGBoost for both the test and external set. A threshold 7.55 (*pK_i_* value) was used for the Metz data as suggested by Pahikkala et al.^12^

**Figure 4:**
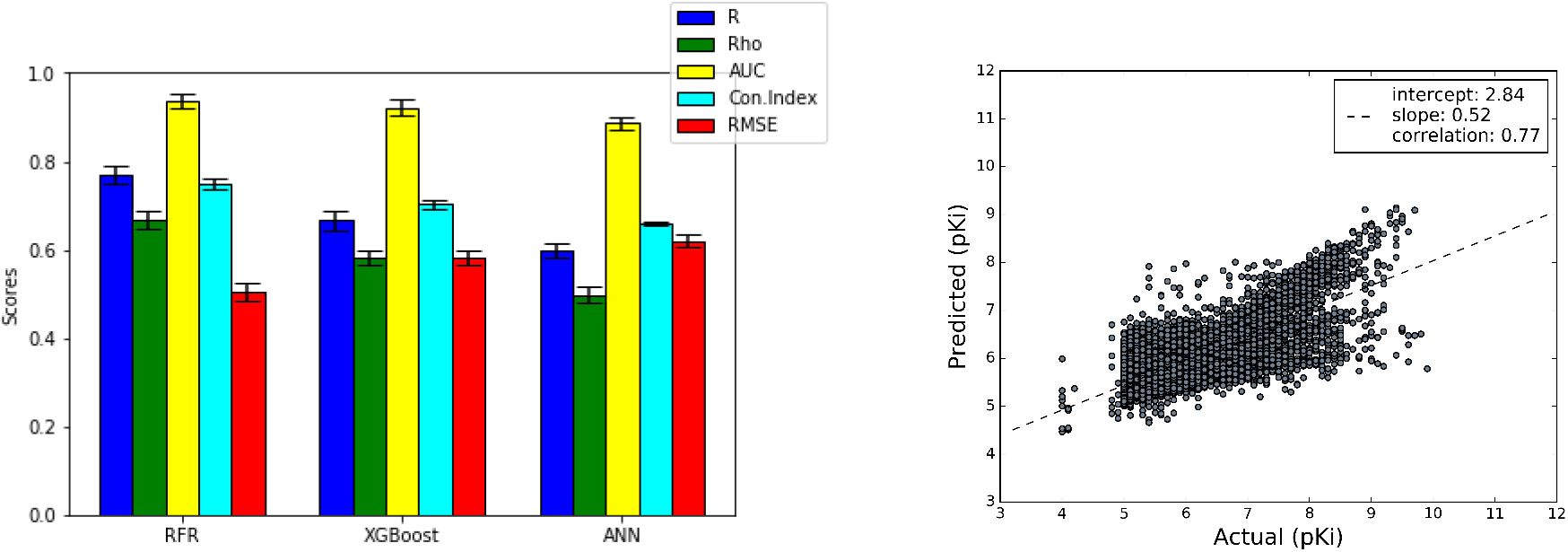
(a) Comparison of models on Metz dataset(b) Predictions from Random Forest against actual values on Metz dataset

## Implementation

The model is available at https://drugdiscovery.utep.edu/pki/. The web interface takes SMILES patterns and protein sequences as input and provides the predicted results. A docker image is also available at `sirimullalab/kinasepkipred`

## Conclusion

A predictive model to estimate the inhibitor constant(*pK_i_*) values of ligands against kinases was developed. Three different ML algorithms were employed and evaluated using a blind test set as well as an independent external set. Out of three models [random forest regressor, extreme gradient boosting (XGBoost), and artificial neural network (ANN)], RFR gave superior results for our drug-target *pK_i_* prediction. Also, a performance test on a grid search and random search was performed, and our results indicated that both methods yielded similar performance. Our best model, which is based on RFR, yielded Pearson R values of 0.887(0.881, 0.893) and 0.769(0.759, 0.779) with 95% confindence interval on the test and Metz datasets, respectively.

## Acknowledgement

We thank Drs. Mahesh Narayan, Amy Wagler, and Gabriel A. Frietze at UTEP for helpful discussions and Dr. Michael Scott Long for reading. We thank the High-Performance Computing Center at UTEP for assistance in using the Chanti cluster and the Texas Advanced Computing Center (TACC) at The UT Austin for providing HPC resources that have contributed to the research results reported within this paper.

## Funding

G.K was supported by the UTEP Computational Science Program. M.H and computational resources were supported by Dr. Sirimulla’s startup fund from the UTEP School of Pharmacy.

## Notes

https://github.com/sirimullalab/KinasepKipred

## References

(1) Pauwels, E.; Stoven, V.; Yamanishi, Y. BMC Bioinformatics 2011, 12, 169.

(2) Moriaud, F.; Richard, S. B.; Adcock, S. A.; Chanas-Martin, L.; Surgand, J.-S.; Ben Jel-loul, M.; Delfaud, F. Briefings in Bioinformatics 2011, 12, 336–340.

(3) Whitebread, S.; Hamon, J.; Bojanic, D.; Urban, L. Drug Discovery Today 2005, 10, 1421 – 1433.

(4) Hopkins, A. L. Nature 2009, 462, 167 EP –.

(5) Anderson, A. C. Chemistry & Biology 2003, 10, 787 – 797.

(6) Kundu, I.; Paul, G.; Banerjee, R. RSC Adv. 2018, 8, 12127–12137.

(7) Yu, L.; Xu, L.; Xu, M.; Wan, B.; Yu, L.; Huang, Q. Molecular Simulation 2011, 37, 1143–1150.

(8) Fabbro, D. Molecular Pharmacology 2015, 87, 766–775.

(9) Gross, S.; Rahal, R.; Stransky, N.; Lengauer, C.; Hoeflich, K. P. The Journal of Clinical Investigation 2015, 125, 1780–1789.

(10) Yamanishi, Y.; Araki, M.; Gutteridge, A.; Honda, W.; Kanehisa, M. Bioinformatics (Oxford, England) 2008, 24, i232–i240, 18586719[pmid].

(11) Li, Q.; Lai, L. BMC Bioinformatics 2007, 8, 353–353, 17883836[pmid].

(12) Pahikkala, T.; Airola, A.; PietilÃ’, S.; Shakyawar, S.; Szwajda, A.; Tang, J.; Aittokallio, T. Briefings in Bioinformatics 2014, 16, 325–337.

(13) He, T.; Heidemeyer, M.; Ban, F.; Cherkasov, A.; Ester, M. J Cheminform 2017, 9, 24–24, 29086119[pmid].

(14) ÃŨztÃĳrk, H.; Ozkirimli, E.; Ozgur, A. Bioinformatics 2018, 34.

(15) Davis, M. I.; Hunt, J. P.; Herrgard, S.; Ciceri, P.; Wodicka, L. M.; Pallares, G.; Hocker, M.; Treiber, D. K.; Zarrinkar, P. P. Nature Biotechnology 2011, 29, 1046 EP –.

(16) Metz, J. T.; Johnson, E. F.; Soni, N. B.; Merta, P. J.; Kifle, L.; Hajduk, P. J. Nature Chemical Biology 2011, 7, 200 EP –.

(17) Tang, J. et al. Cell Chemical Biology 2018, 25, 224 – 229.e2.

(18) Cao, D.-S.; Xu, Q.-S.; Liang, Y.-Z. Bioinformatics 2013, 29, 960–962.

(19) Sud, M. Journal of Chemical Information and Modeling 2016, 56, 2292–2297.

(20) `relax`mciteBstWouldAddEndPuncttrue.

(21) McGregor, M. J.; Muskal, S. M. Journal of Chemical Information and Computer Sciences 1999, 39, 569–574.

(22) Horvath, D. RSC Publishing. 44–75. 2008,

(23) Ewing, T.; Baber, J. C.; Feher, M. Journal of Chemical Information and Modeling 2006, 46, 2423–2431.

(24) Watson, P. Journal of Chemical Information and Modeling 2008, 48, 166–178.

(25) Bonachéra, F.; Parent, B.; Barbosa, F.; Froloff, N.; Horvath, D. Journal of Chemical Information and Modeling 2006, 46, 2457–2477.

(26) Pedregosa, F. et al. Journal of Machine Learning Research 2011, 12, 2825–2830.

(27) Bergstra, J.; Bengio, Y. JMLR 2012, 305.

