## Supporting Information for "KinasepKipred: A Predictive Model for Estimating Ligand-Kinase Inhibitor Constant (*pK_i_*)"

October 31, 2019

**2 Evaluation metrics** **3**

**3 Grid search vs random search** **4**

**References** **5**

### 1 Machine learning model development

#### 1.1 Random forest

Random forest [1] is an ensemble technique capable of performing both regression and classification tasks. It uses multiple decision trees and trains each decision tree based on technique called bootstrap aggregation, which is also known as bagging. It creates decision trees on randomly selected data samples, obtains predictions from each tree, and selects the best solution by means of voting [2]. It is widely used for a variety of applications, such as recommendation engines, image classification and feature selection. We used three hyperparameters, (number of estimators, min sample split and maximum features) to optimize the models. Our best model was obtained with the hyperparameter combination of "number_estimators": 600, "min_samples_split": 2, and "max_features": "auto".

#### 1.2 Extreme gradient boosting

Extreme gradient boosting (XGBoost) [3] is an implementation of gradient boosted decision trees designed for speed and performance. The utility of this algorithm lies in its scalability, which drives fast learning through parallel and distributed computing and offers efficient memory use. We used the random search method to tune the hyperparameters in XGBoost within a given parameter space. Among many hyperparamters, n_estimators, objective, colsample_bytree, learning_rate, max_depth, and alpha are found effective for tuning the model. Our best XGsboost model was developed with a combination of "n_estimators": 500, "objective": "reg_linear", "colsample_bytree": 0.3, "learning_rate": 0.1, "max_depth": 5 and "alpha": 10.

#### 1.3 Artificial neural network

Artificial neural networks (ANNs) [4] are the brain-inspired systems which are intended to replicate the way that we humans learn. They are comprised of artificial neurons, also known as nodes. An ANN usually consists of input layers, output layers, and hidden layers most of the time. Input layers, also known as synapses are passed to the neurons in the hidden layers. Synapses are assigned with weights. Weights play a crucial role to select important signals for the output. Signals that reach into the neurons are summed to form a weighted sum. This weighted sum is applied to the activation function. Based on the activation function neuron will either pass on the signal or not. We used an input layer with 32 nodes, and 5 hidden layers as 32, 64, 128,128, 64, 32, and an output layer. As an activation function, we used a rectified linear unit (ReLU) [5]. The final layer is activated by a linear function to predict the output. Since the model was regression based, we used mean squared error (MSE) as the 'loss function' and 'adam' as an optimizer. The learning was completed in 100 epochs with batch size 25.

#### 2 Evaluation metrics

**Concordance index:** Since the binding affinities in drug target interactions are continuous values, one of the evaluation metrics used in this study was the concordance Index (Con. Index) ([6]). Con. Index measures the probability of two randomly drawn drug-target pairs with different label values are in the correct order. In other words, the prediction for the larger affinity is larger than the prediction of the smaller affinity value. If $d_i$, $d_j$, $x_i$ and $x_j$ are the predicted and the actual affinity values in the case of larger and smaller binding affinities respectively, Con. Index can be obtained as:

$$Con \cdot Index = \frac{1}{Z} \sum_{x_i > x_j} h(d_i - d_j) \tag{1}$$

where Z is the normalization constant, and h(u) is the step function. h(u) is 1.0, 0.5 and 0.0 for $u > 0$, $u = 0$ and $u < 0$ respectively. The Con. Index values lies in between 0.5 and 1.0. The value 0.5 corresponds to a random prediction, and as 1.0 for a perfect prediction.

**Root mean square Error(RMSE):** Define $B_a$ and $B_p$ as the actual and predicted binding affinities. The error can be calculated as:

$$Error(E_i) = B_{a_{(i)}} - B_{p_{(i)}} \tag{2}$$

MSE is calculated as the mean of the differences of the actual binding affinity values and the calculated binding affinity values.

$$MSE = \frac{1}{N} \sum_{i=1}^{N} E_i^2 \tag{3}$$

The RMSE is obtained as:

$$RMSE = \sqrt{MSE} \tag{4}$$

**Pearson's correlation coefficient (R):**

$$\overline{B_a} = \frac{1}{N} \sum_{i=1}^{N} B_{a_{(i)}} \tag{5}$$

$$\overline{B_p} = \frac{1}{N} \sum_{i=1}^{N} B_{p_{(i)}} \tag{6}$$

$$R = \frac{\sum_{i=1}^{N} (B_{a_{(i)}} - \overline{B_a})(B_{p_{(i)}} - \overline{B_p})}{\sqrt{\sum_{i=1}^{N} (B_{a_{(i)}} - \overline{B_a})^2 \sum_{i=1}^{N} (B_{p_{(i)}} - \overline{B_p})^2}} \tag{7}$$

R measures the linear relationship between the actual and predicted binding affinity scores. It lies in between -1 and 1.

**Area under the curve (AUC):** The area under the Receiver Operating Characteristic

(ROC) curve is generally adopted in binary classification problems. However, it can also be used to measure the regression problems by converting the quantitative values into binary values by selecting thresholds. There are different ways to measure AUC for regression problems. We have measured the average AUC converting the actual compound kinase interaction values (i.e, $K_i$) into binary labels given certain interaction thresholds. The ROC curve was created by plotting the true positive rate (TPR) (i.e., sensitivity) is plotted against the false positive rate (FPR) (i.e., 1 − specificity) for different cut-off points. Accuracy is measured by the area under the ROC curve. An area of 1 (i.e, 100%) represents a perfect test; an area of 0.5 (i.e, 50%) represents a random test.

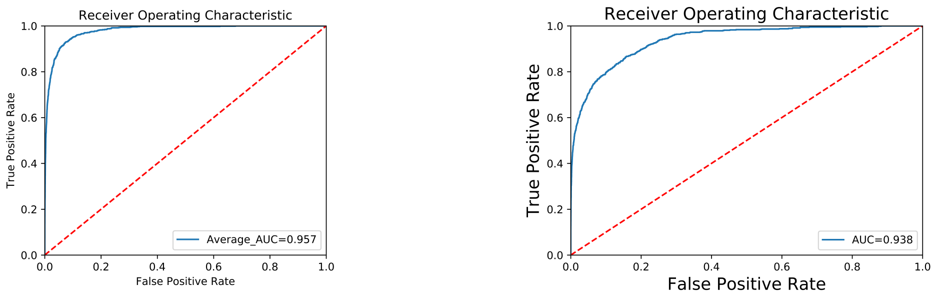

Figure 1: Area under the ROC curve for (a) Test dataset, (b) Metz dataset

### 3 Grid search vs random search

Hyperparameter optimization is considered to be the most challenging aspect of for any model development. The main puropose here is to find a set of hyperparameters which will lead to high precision and accuracy. Random forest was tuned by both grid search, and random search methods, and we compared the performance of the model. In a grid search, a combination of hyperparameters are first used for tuning the model and then the combination which gives the best results results are chosen; whereas in the random search method [7], a distribution of hyperparameters are provided and the values are randomly picked on each iteration to train the model. It was more generally a process of choosing a representative sample of the parameter from the entire space to understand all the data. We used three hyperparameters, namely, n estimators, max features and min sample split for RFR to optimize the model based on both grid search and random search. The model performance was nearly the same for both the methods. However, the model development time was significantly less for random search. This indicates that random search gave similar results in a fraction of the time taken by grid search method. We also observed a change in R and RMSE in accordance with an increasing the number of trees in RFR, yet it was found insignificant. It instead increased model development time. The performance of RFR model by the grid search method and random search method is shown in Table (2) and Table (3 shows the changes in R and RMSE on increasing the number of trees in the RFR model.

Table 1: Hyperparameter tuning in random rorest regressor

| Parameters | Range | Grid Search (best combination) | Random Search (best combination) |
|---|---|---|---|
| n_estimators | [50, 100, 200, 400, 600,800, 1000] | 600 | 600 |
| max_features | [`auto', `sqrt', `log2', None] | 'auto' | None |
| min_sample_split | [2, 5, 10] | 2 | 2 |

Table 2: Comparison of random search and grid Search

| EvaluationMetrics | Random Search | | Grid Search | |
|---|---|---|---|---|
| | Test set | Metz set | Test set | Metz set |
| Pearson | 0.887 | 0.769 | 0.887 | 0.769 |
| Spearman | 0.845 | 0.667 | 0.846 | 0.669 |
| RMSE | 0.475 | 0.504 | 0.475 | 0.504 |
| Concordance Index | 0.850 | 0.740 | 0.854 | 0.749 |
| AUC | 0.946 | 0.937 | 0.957 | 0.938 |

Table 3: Change in R and RMSE on increasing the the number of trees for the test set

| Number of trees | R | RMSE |
|---|---|---|
| 50 | 0.8829 | 0.4826 |
| 100 | 0.8865 | 0.4764 |
| 200 | 0.8871 | 0.4753 |
| 400 | 0.8874 | 0.4748 |
| 600 | 0.8876 | 0.4745 |
| 800 | 0.8871 | 0.4753 |
| 1000 | 0.8873 | 0.4751 |

### References

[1] Leo Breiman. Random forests. *Machine Learning*, 45(1):5–32, Oct 2001.

[2] M.C. Mariani, M. A. Masum Bhuiyan, Sushil Shakyawar, and Osei Tweneboah. Statistical data mining algorithms for the prognosis of diabetes and autism. *Hawaii University International Conferences, Hawaii, June 5-7*, 2019.

[3] Tianqi Chen and Carlos Guestrin. XGBoost: A scalable tree boosting system. In *Proceedings of the 22nd ACM SIGKDD International Conference on Knowledge Discovery and Data Mining*, KDD '16, pages 785–794, New York, NY, USA, 2016. ACM.

[4] Warren S McCulloch and Walter Pitts. A logical calculus of the ideas immanent in nervous activity. *The bulletin of mathematical biophysics*, 5(4):115–133, 1943.

[5] Vinod Nair and Geoffrey E. Hinton. Rectified linear units improve restricted boltzmann machines. In *Proceedings of the 27th International Conference on International Conference on Machine Learning*, ICML'10, pages 807–814, USA, 2010. Omnipress.

[6] Tapio Pahikkala, Antti Airola, Sami Pietilä, Sushil Shakyawar, Agnieszka Szwajda, Jing Tang, and Tero Aittokallio. Toward more realistic drug-target interaction predictions. *Brief Bioinform*, 16(2):325–337, Mar 2015. 24723570[pmid].

[7] James Bergstra and Yoshua Bengio. Random search for hyper-parameter optimization. *JMLR*, page 305, 2012.
